# Phosphorylation of Cyclophilin-D is Not Required for Regulation of The Mitochondrial Permeability Transition Pore by GSK3β

**DOI:** 10.64898/2026.01.15.699680

**Authors:** Linda Alex, Paula J. Klutho, Lihui Song, Manuel Gutiérrez-Aguilar, Christopher P. Baines

## Abstract

Genetic inhibition of cyclophilin D (CypD) delays the opening of the mitochondrial permeability transition pore (MPTP) and therefore reduces necrotic cell death. Elucidation of factors that impact CypD activity is therefore key to understanding the regulation of MPTP opening. Glycogen synthase kinase-3β (GSK3β) is a serine/threonine kinase that has been shown to modulate MPTP and cell death, potentially through phosphorylation of CypD. Therefore, we hypothesized that the mitochondrial fraction of GSK3β directly phosphorylates CypD and promotes opening of MPTP. Overexpression of full length GSK3β in mouse embryonic fibroblasts sensitized the MPTP and exacerbated oxidative stress-induced necrosis. In contrast, genetic inhibition of GSK3β protected against oxidant-induced cytotoxicity but did not affect the MPTP. Recombinant GSK3β could directly bind to and phosphorylate recombinant CypD. Mass spectrometry revealed several putative GSK3β phosphorylation sites on CypD. However, mutation of these sites did not affect the peptidyl prolyl isomerase activity of CypD and reconstitution of these phosphomutants in CypD-deficient cells increased MPTP sensitivity and oxidative-induced cell death to the same extent as wild-type CypD. Further, targeted overexpression of either wild-type or kinase-inactive GSK3β in the mitochondrial matrix did not impact MPTP or cell death. Moreover, while proteinase-K digestion of cardiac mitochondria showed a significant amount of GSK3β in the mitochondria, it was not localized to the matrix. Finally, overexpression of GSK3β was still able to increase MPTP sensitivity and oxidative stress-induced death in CypD-null cells. Taken together, these data indicate that, while GSK3β can modulate MPTP, this appears to be independent of GSK3β’s interaction with, or phosphorylation of CypD.

## Introduction

Mitochondrial permeability transition (MPT) mediates a regulated form of necrotic cell death implicated in multiple pathological states like cardiac ischemic-reperfusion injury [1], liver toxicity [2], lysosomal storage diseases [3], neurodegenerative disorders [4] and muscle myopathies [5]. On a molecular basis, MPT is mediated by the opening of the MPT pore (MPTP), induced by elevated Ca^2+^ levels in the mitochondrial matrix and potentiated by ROS upsurge, depleted ATP levels and increased phosphate ion concentration [1,2,3,4,5]. Physiological opening of MPTP, also called “flickering MPTP”, aids cells in the tightly controlled flux of Ca^2+^ across mitochondrial membranes [6,7]^7^. However, upon injury, the MPTP can remain persistently opened, allowing flux of solutes up to 1500 Da, resulting in the dissipation of transmembrane potential (ΔΨ) which inhibits ATP synthesis [1,2,3,4,5,6,7]. Mitochondrial swelling ultimately results in the rupture of mitochondrial membranes and finally leads to necrotic cell death.

MPTP was initially hypothesized to comprise the voltage-dependent anion channel (VDAC) on the outer mitochondrial membrane, adenine nucleotide translocase (ANT) in the inner mitochondrial membrane and cyclophilin D (CypD) in the mitochondrial matrix [1,2,3,4,5,6,7]. However, classical genetic knockout mouse model studies showed that MPT occurs even in the absence of these putative components indicating that they may not constitute the core structure of the pore [8,9,10]. Interestingly, CypD-encoding gene abrogation or pharmacological inhibition delays, but does not inhibit, MPT and reduces cell death, indicating that CypD is not a channel-forming component of the MPTP but rather is a pore regulator [11,12,13,14,15]. However, little is known regarding the molecular effectors that directly regulate CypD to modulate MPT.

CypD, a mitochondrial matrix protein, has peptidyl-prolyl isomerase (PPIase) activity that is required for its ability to facilitate MPTP opening [11,12,16]. Regulation of CypD activity appears to be controlled primarily through post-translational modifications such as phosphorylation [17,18,19], acetylation [20,21], and S-nitrosylation [22]. Glycogen synthase kinase (GSK)-3β is a candidate serine/threonine kinase that’s been shown to stimulate MPT and cell death. Inhibition of GSK3β delays MPT and protects cells from death induced by a variety of stimuli [17,18,23,24]. Co-immunoprecipitation studies have also shown that GSK3β may localize to the mitochondrial matrix and physically interact with CypD [17,24], and manipulation of GSK3β affects CypD phosphorylation status [17,18,25,26]. However, a direct validation of how GSK3β-CypD phosphoregulation impacts MPT is still lacking.

In the present study, we hypothesized that mitochondrial GSK3β directly phosphorylates CypD to increase its activity and therefore enhance opening of MPTP and cell death. Mass spectrometry and bioinformatics tools showed that GSK3β could phosphorylate CypD *in vitro* at multiple serine/threonine residues. So, we developed CypD-encoding recombinant adenoviruses mutated at these residues to either aspartate to mimic phosphorylation, or alanine to block the phosphorylation. However, contrary to our expectations, we found that GSK3β-mediated MPT occurs independently of CypD phosphorylation. Further, overexpression of mitochondrially-targeted wild-type (WT) GSK3β as well as dominant negative (DN) GSK3β did not impact MPT suggesting that the pro-MPT effects of GSK3β may be independent of the mitochondrial pool. Also, corroborating to this, we found that full length GSK3β sensitizes cells to MPT even in the absence of CypD, thereby suggesting an alternative pathway of GSK3β-sensitization of the MPTP independent of CypD.

## Materials and Methods

### Materials

Dulbecco’s Modified Eagles Medium (DMEM), pencillin/streptomycin, trypsin/EDTA, Calcium Green-5N, BL21 cells, Lipofectamine RNAiMAX, Halt protease/phosphatase inhibitor, and HisPur Co^2+^ affinity purification columns were from Thermo Fisher Scientific. Fetal Bovine Serum (FBS) was from Atlanta Biologicals. The pAcGFP1-mito vector was from Takara. The pGEX-4T1 vector and Enhanced Chemifluorescence (ECF) reagent were from GE Healthcare Life Sciences. The pCMV-Tag1 vector and AdEasy-XL kit were from Agilent Technologies. Recombinant His-tagged human CypD was from ProSpec. Recombinant active GST-GSK3β and Ponceau-S protein stain were from Millipore-Sigma. Kinase buffer and ATP were from Cell Signaling. All other reagents were from either Millipore-Sigma or Thermo Fisher Scientific.

### Animal Use and Approval

Isolation of mouse embryonic fibroblasts (MEFs) and harvesting of mouse hearts were conducted in accordance with the National Institutes of Health Guidelines for the Care and Use of Laboratory Animals. All procedures including methods of euthanasia were submitted to and approved by the Animal Care and Use Committee of the University of Missouri.

### Cell Culture

The CypD-deficient (*Ppif*^−/−^) mice have been described previously [12]. Pregnant dams were euthanized at embryonic day 15.5-16.5 and the embryos excised. The head, appendages, internal organs were removed (the tail was used for genotyping), and the remaining tissue digested in 0.05% trypsin/1mM EDTA for 30min at 37°C. The suspensions were then plated in DMEM containing 10% FBS, and 100U/mL penicillin, and 0.1mg/mL streptomycin. Cells at passages 2-5 were used for experiments. Each isolate was considered one biological replicate.

### Adenoviruses and siRNA

Replication-deficient adenoviruses encoding wild-type GSK3β (WT-GSK3β and mutant GSK3β kinase-inactivating mutation (K85A, DN-3β) were a generous gift from Dr. Junichi Sadoshima at Rutgers University. The adenovirus for β-galactosidase (βGal) has been previously described [9,12]. To generate the mitochondrial-targeted viruses, we obtained clones for HA-tagged human wild-type and K85A GSK3β with HA-tags as a gift from Dr. James Woodgett at (Addgene plasmid #14753 http://n2t.net/addgene:147553; RRID:Addgene_14753 and #14755 http://n2t.net/addgene:14755; RRID:Addgene_14755, respectively). The cDNAs were subcloned into the *Bam*H1 and *Not*I sites of pAcGFP1-mito, placing them distal to and in frame with the human COXVIII mitochondrial targeting sequence and replacing the AcGFP sequence to generate mitochondrial targeted WT and DN-GSK3β (miWT-3β and miDN-3β). Having confirmed mitochondrial localization of the constructs in HEK293 cells, the cDNAs were then subcloned into the *Kpn*I and *Xho*I sites of pShuttle-CMV. For the CypD constructs, mouse CypD was subloned into the *Not*I and *Bam*H1 sites of pCMV-Tag1 to generate CypD with a C-terminal FLAG tag. Phosphorylation mutants of CypD-FLAG were then generated using the QuikChange-II mutation kit. The different CypD-FLAG cDNAs were then subcloned into the *Kpn*I and *Xho*I sites of pShuttle-CMV. Adenoviruses encoding the mito-targeted GSK3β and CypD-FLAG constructs were then generated using the AdEasy-XL kit. MEFs were infected 48hrs before experimentation with adenovirus at a multiplicity of infection of 100-200.

For the knockdown experiments, MEFs were transfected with a pool of 4 mouse-specific GSK3β targeting siRNAs (M-053232-01, Dharmacon: #1 5’-gcagaaguguacaaacgaa-3’, #2 5’-gggcugugcguuuaaugua-3’, #3 5’-cuacgaagcaucucuuuac-3’, #4 5’-ggagaauggcuucacgaau-3’) at a concentration of 100nM using Lipofectamine RNAiMax for 48 hrs before experimentation. A pool of 4 non-targeting siRNAs was used as a control (D-001206-14, Dharmacon: #1 5’-uaaggcuaugaagagauac-3’, #2 5’-auguauuggccuguauuag-3’, #3 5’-augaacgugaauugcucaa-3’, #4 5’-ugguuuacaugucgacuaa-3’).

### Calcium Retention Capacity and Cell Death Assays

For assessment of calcium retention capacity (CRC), infected/transfected MEFs were trypsinized, counted, washed with PBS and then resuspended in CRC buffer (120mM KCl, 10mM Tris at pH 7.4, 1mM KH_2_PO_4_, 20μM EDTA) at a concentration of 4×10^6^ cells/mL. For each measurement 1×10^6^ cells, 5mM succinate, 1μM Calcium Green-5N and 8μM digitonin were loaded into a cuvette to a total volume of 1mL. The cuvette was placed in a fluorometer and exposed to 500nm light. After two minutes, pulses of 2.5μM CaCl_2_ were added every minute until an increase in fluorescence was detected at 530nm consistent with MPT pore opening. To induce cell death, infected/transfected MEFs were treated with increasing concentrations of H_2_O_2_ for 4hrs. At the end of the protocol cells were co-stained with the vital dye Sytox Green (0.15μM) (Invitrogen) to label dead cells and bis-benzimide (10μg/mL) to label all cells for 15min in PBS. Images were captured on an inverted fluorescence microscope (Olympus IX51) and analyzed using ImageJ (NIH).

### Mitochondrial Isolation and Assays

Cardiac mitochondria were prepared from wild-type mice by differential centrifugation in sucrose-based medium. Mice were euthanized and the hearts excised and rinsed in ice-cold PBS. The hearts were homogenized using a Dounce in a sucrose buffer (250mM sucrose, 10mM Tris pH7.4, 1mM EDTA). The crude homogenates were centrifuged at 1,000*g* for 5min at 4°C to pellet nuclei and tissue debris. The supernatant was removed and spun at 10,000*g* at 4°C for 10min. The mitochondrial pellet was then washed twice in sucrose buffer while the supernatant was subjected to further centrifugation at 100,000g for 60min at 4°C to generate the cytosolic fraction. To separate integral versus associated membrane proteins, mitochondria were incubated in 100mM Na_2_CO_3_ at pH11.0, for 30min at 4°C. The insoluble membrane proteins were then pelleted by centrifugation at 100,000*g* for 30 min at 4°C. Both the soluble and membrane fractions were then subjected to Western blotting. For the digestion assays, purified mitochondria were incubated with increasing concentrations of proteinase-K (2–200μg/ml) for 20 min on ice. The digestion was then stopped by the addition of 1mM PMSF and the mitochondria subjected to Western blotting.

### Protein Interaction Assays

To assess direct interactions, 1μg of recombinant His-tagged CypD was incubated with increasing amounts (0.2-2.0μg) of recombinant GST-GSK3β in 1mL of binding buffer (1% NP-40 in PBS) overnight at 4°C. The mixture was then spun through a Co^2+^-conjugated agarose column to bind the His-tagged CypD. After washing 3 times with binding buffer, the complexes were eluted (300mM NaCl, 150mM imidazole) and subjected to Western blotting. For immunoprecipitation from cells, MEFs were infected with β-galactosidase, WT-GSK3β, and/or CypD-FLAG for 48hrs. Cells were then lysed for 30min on ice in 1mL of immunoprecipitation buffer (150mM NaCl, 20mM Tris pH7.4, 1mM EDTA, 10% glycerol, 0.2% NP40, protease/phosphatase inhibitor). Lysates were clarified by centrifuging at 17,000*g* for 20min at 4°C and protein concentration determined by the Bradford assay. One mg of each sample was then incubated overnight with 25μL of anti-FLAG-conjugated agarose (A2220, Sigma) in a final volume of 1mL. After washing 3 times with immunoprecipitation buffer the beads were subjected to Western blotting.

### Proteomic Analyses

For intact mass analysis, a 5μL aliquot of sample was subjected to a short LC-MS run using the following parameters: acquired mass range 300-2500m/z; 0.63 spectra/second; fragmentor at 200V. Gradient was as follows: initial conditions (for trap load) was 3% B (A: 0.1% formic acid in water; B: 99.9% acetonitrile, 0.1% FA); rapid ramp to 20% B over 1min; hold at 20%B for 3min; gradient 20-60% B over 6min; ramp to 90%B over 1min; hold at 90% B for 4min; ramp back to initial conditions over 30sec and hold at 3%B for 3min prior to loading next sample. Samples (0.5μL - 500fmol) were loaded in sequence as follows: sample; blank (1% FA). Data were then examined using the software provided with the instrument.

For the phospho-mapping, data were acquired by LC-MS on the LTQ Orbitrap XL using an MSA (multistage activation) method. This method looks specifically for “neutral losses” of the phosphate from peptides, which typically occur with (Ser/Thr) phospho-peptides. Once a neutral loss is found, a second round of MSMS is done on the “neutral loss peak”. These two spectra are then combined into a single spectrum for database searching. To this end, the remainder of the sample not used for intact mass analysis was lyophilized sample and then resuspended in 10μL of 6M Urea/100mM HEPES pH8.0. Following reduction and alkylation of cysteines, the proteins were digested with trypsin overnight “in-solution”. The reaction was then acidified, lyophilized again, and resuspended in 10μL of 1% formic acid. A portion of the digest (5 μL) was loaded directly onto a 25 cm long, 150 μm inner diameter, pulled-needle analytical column packed with Magic C18 reversed phase resin (Michrom Bioresources). Peptides were separated and eluted from the analytical column with a gradient of acetonitrile as follows: initial conditions (during trap load) and for the first 2min was 0% B (A: 0.1% formic acid in water; B: 99.9% acetonitrile, 0.1% FA), followed by a gradient from 0 to 40%B over 90min, ramp to 90% B over 1min, hold at 90% B for 15min, ramp to 5%B over 1min, hold at 5%B for 6 min prior to loading next sample.

The Proxeon Easy nLC system is attached to an LTQ Orbitrap XL mass spectrometer. Following a high-resolution (30,000res, profile) FTMS scan of the eluting peptides (300-2000m/z range), each cycle, the 9 most abundant peptides (reject porcine trypsin autolysis ions, reject +1 ions, >2000 counts) were subjected to iontrap CID peptide fragmentation (NCE of 35%, centroid, isolation width of 2). Dynamic exclusion was enabled with repeat count of 1, repeat duration of 30sec, exclusion list of 500, and exclusion duration of 120sec. Data across a total of 130 minutes of elution were collected. Raw data were copied to the Sorcerer and searched against the IPI-Human and SwissProt-all species databases with 25ppm mass error tolerance, monoisotopic mass, and oxidized Met and phospho STY as variable mods. The Sorcerer output file was imported into the Scaffold software V3.4.8 and examined for hits.

### CypD Activity Assay

The wild-type and phospho-CypD constructs that were generated for the adenoviruses were subcloned into the *Eco*R1 and *Xho*I sites of pGEX-4T1 downstream and in-frame of the GST cDNA. The cDNA were transfected in to BL21 cells and protein expression induced with IPTG. The bacteria were then lysed and the GST proteins purified using glutathione-conjugated sepharose beads. After extensive washing the proteins were eluted with excess GSH, dialyzed and purity determined by SDS-PAGE followed by Coomassie staining. PPIase activity was determined as previously described [22]. Briefly, 1μg of each protein was added to 800μL of assay buffer (100mM NaCl, 50mM HEPES pH8.0) containing 0.25mg/mL chymotrypsin, and 300nM of the substrate N-succinyl-Ala-Ala-Pro-Phe-*p*-nitroanilide. The enzymatic activity of CypD converts the substrate from a *cis* to *trans* form, thereby allowing chymotrypsin to cleave off the *p*-nitroanilide. The free *p*-nitroanilide can then be measured in a spectrophotometer at 410nm. GST alone was used as a negative control for the GST-CypD proteins. Cyclosporin-A (CsA, 1μM) was used to chemically inhibit CypD activity. CypD-driven isomerization rates were calculated using a non-linear regression and expressed as the observed rate (K) minus the spontaneous rate (K0).

### Protein Modeling

Cyclophilin D was modeled with PyMol (The PyMol Molecular Graphics System, Version 2.3 Schrödinger, LLC) using PDB structure 3QYU.

### Western Blotting

Cells and mitochondrial pellets were homogenized in buffer (150mM NaCl, 10mM Tris pH7.4, 1mM EDTA, 1% Triton-X100, protease/phosphatase inhibitor) and then centrifuged at 17,000*g* for 10min at 4°C to remove debris. Protein concentration was determined by the Bradford assay. SDS loading buffer (final concentration 2% SDS, 8% Glycerol, 4% b-mercaptoethanol, 0.01% Bromophenol Blue, 50mM Tris pH7.0) was added to the homogenates, immunoprecipitates, or recombinant proteins and the samples resolved on 10% acrylamide gels by SDS-PAGE. The proteins were then transferred onto PVDF membranes. After blocking in 10% non-fat milk in TBS-T, primary antibodies: AIF (BD Biosciences, #551429, 1:500), CypD (Abcam, ab110324, 1:1000), GAPDH (Millipore-Sigma, MAB374, 1:1000), GSK3b (Cell Signaling, 9315S, 1:1000), GST (Abcam, 1:1000) FLAG (Sigma, F7425, 1:2000), HA (Millipore-Sigma, 04-902, 1:1000), HK2 (Abcam, ab78259, 1:1000), LDH (Abcam, ab227198, 1:1000), Myc (Cell Signaling, 2276S, 1:1000), OXPHOS antibody cocktail (Abcam, ab110413, 1:1000), phosphoserine/phosphothreonine mix (Millipore-Sigma, AB1603 and AB1607, 1:500), and VDAC (Abcam, ab14734 1:1000) were applied to the membranes overnight at 4°C in blocking buffer. After washing in TBS-T, membranes were incubated with the appropriate alkaline phosphatase-linked secondary antibody (Cell Signaling, 1:1000) and visualized by enhanced chemifluorescence.

### Immunocytochemistry

MEFs were cultured in chamber slides and infected with the HA-tagged miWT-GSK3β and miDN-GSK3β adenoviruses for 48hrs. The MEFs were then washed with PBS, fixed with 4% paraformaldehyde, and permeabilized/blocked with blocking solution (PBS containing 1% bovine serum albumin, 0.1% cold water fish skin gelatin, 0.1% Tween-20) for 1hr at RT. The slides were then incubated overnight at 4°C with anti-HA (Millipore-Sigma, 04-902, 1:500) and anti-CypD (Abcam, ab110324, 1:500) antibodies in blocking buffer. After washing 3 times with PBS/0.1% NP-40, cells were incubated with the appropriate fluorophore-conjugated secondary antibody (Alexa, Thermo Scientific, 1:500) for 2hrs at RT. The cells were washed 3 more times and imaged using an inverted fluorescence microscope.

### Statistical Analyses

Student’s unpaired *t*-test was used to determine statistical significance between two groups. A One-Way ANOVA followed by Scheffé’s post-hoc test was used for comparison of multiple groups. For experiments where both genetic manipulation and H_2_O_2_ concentration were factors, a Two-Way ANOVA was employed followed by Scheffé’s post-hoc test. A p value <0.05 was considered statistically significant.

## Results

### Overexpression of GSK3β sensitizes MEFs to MPT and oxidative stress-induced cell death

We first wanted to assess the effects of GSK3β overexpression on MPT sensitivity and cell death. To this end, MEFs were infected with adenoviruses encoding either β-galactosidase (βGal, control) or wild-type GSK3β (WT-GSK3β) for 48 hrs (**Figure 1A**). Treatment with increasing concentrations of H_2_O_2_ for 4 hrs elicited a concentration-dependent increase in Sytox positivity in the βGal-infected MEFs (**Figure 1B**), an effect that was significantly exacerbated in the WT-GSK3β overexpressing cells (**Figure 1B**). To assess MPT, we measured Ca^2+^ retention capacity (CRC) in the infected MEFs. Representative CRC traces are depicted in **Figure 1C**. Calcium (2.5 μM)-induced spikes in fluorescence were observed every minute followed by a decrease in fluorescence as the mitochondria took up the Ca^2+^. Subsequent opening of the MPT pore, resulted in a large amplitude increase in fluorescence as the Ca^2+^ was released. The number of Ca^2+^ spikes was then used to calculate the CRC of each sample. Overexpression of WT-GSK3β resulted in a significant reduction in CRC when compared to βGal controls (**Figure 1C,D**), indicative of an increased MPT sensitivity. These data indicate that overexpression of exogenous GSK3β can enhance MPT responsiveness to Ca^2+^ and increase oxidative stress-induced cell death.

**Figure 1:**
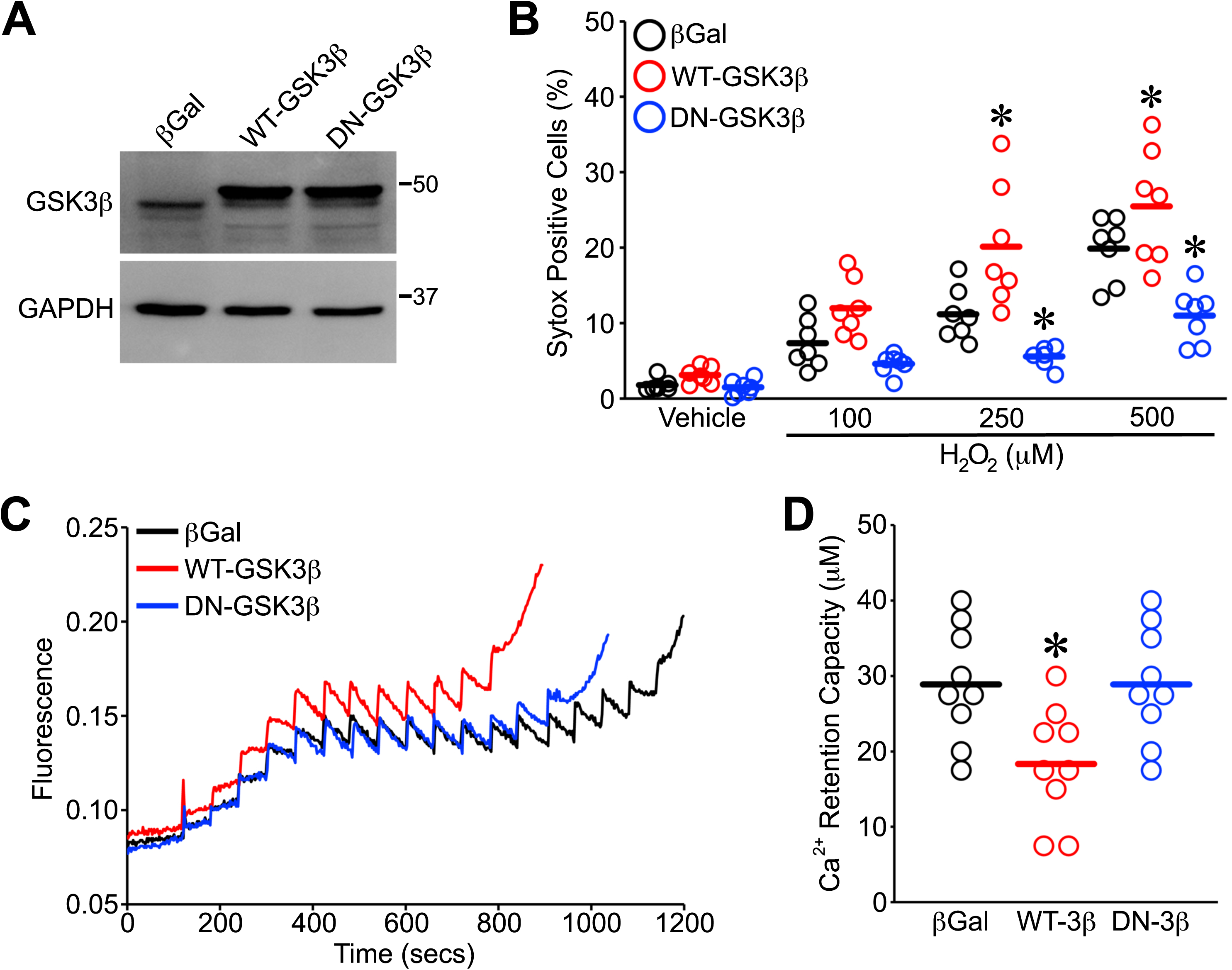
Overexpression of GSK3β sensitizes MEFs to MPT and oxidative stress-induced cell death. Wild-type MEFs were transfected with adenovirus encoding βGal (control), wild-type (WT) or kinase-active (K85R, DN) GSK3β for 48hrs. **A**) Western blotting for GSK3β demonstrating equivalent expression of the WT-GSK3β and DN-GSK3β. GAPDH was used as a loading control. **B**) βGal, WT-GSK3β and DN-GSK3β-infected MEFs were treated with H_2_O_2_ (100, 250 or 500μM) for 4hrs and cell death was measured by Sytox green staining. **C**) Representative traces of Ca^2+^ retention capacity (CRC) measured using Calcium Green-5N in digitonin-permeabilized βGal, WT-GSK3β and DN-GSK3β-infected MEFs. Cells were treated with 2.5 μM Ca^2+^ boluses every minute until the peak of fluorescence indicative of MPT was observed. **D**) Quantification of CRC data in the βGal, WT-GSK3β and DN-GSK3β-infected MEFs. Each individual point represents one independent cell isolate. Bar represents the mean. \**P*<0.05 vs. βGal.

### Inhibition of GSK3β protects MEFs against oxidative stress but does not affect MPT

We next tested the effects of GSK3β inhibition on MPT and cell death using two different approaches. In contrast to WT-GSK3β, overexpression of kinase-inactive GSK3β (DN-GSK3β) (**Figure 1A**) significantly protected MEFs against cell death induced by increasing H_2_O_2_ concentrations (**Figure 1B**). However, this protective effect was independent of any changes in CRC (**Figure 1C,D**). We also depleted cells of GSK3β using siRNA (**Figure 2A**). Similar to the dominant-negative intervention, reduction of endogenous GSK3β provided significant protection against H_2_O_2_-induced cytotoxicity when compared to MEFs transfected with a non-targeting siRNA (**Figure 2B**). Again, however, this was not associated with any changes in CRC (**Figure 2C,D**). Thus, as previously shown, inhibition of GSK3β protects against oxidant-induced death. However, in our hands this appears to be independent of any changes in MPT responsiveness.

**Figure 2:**
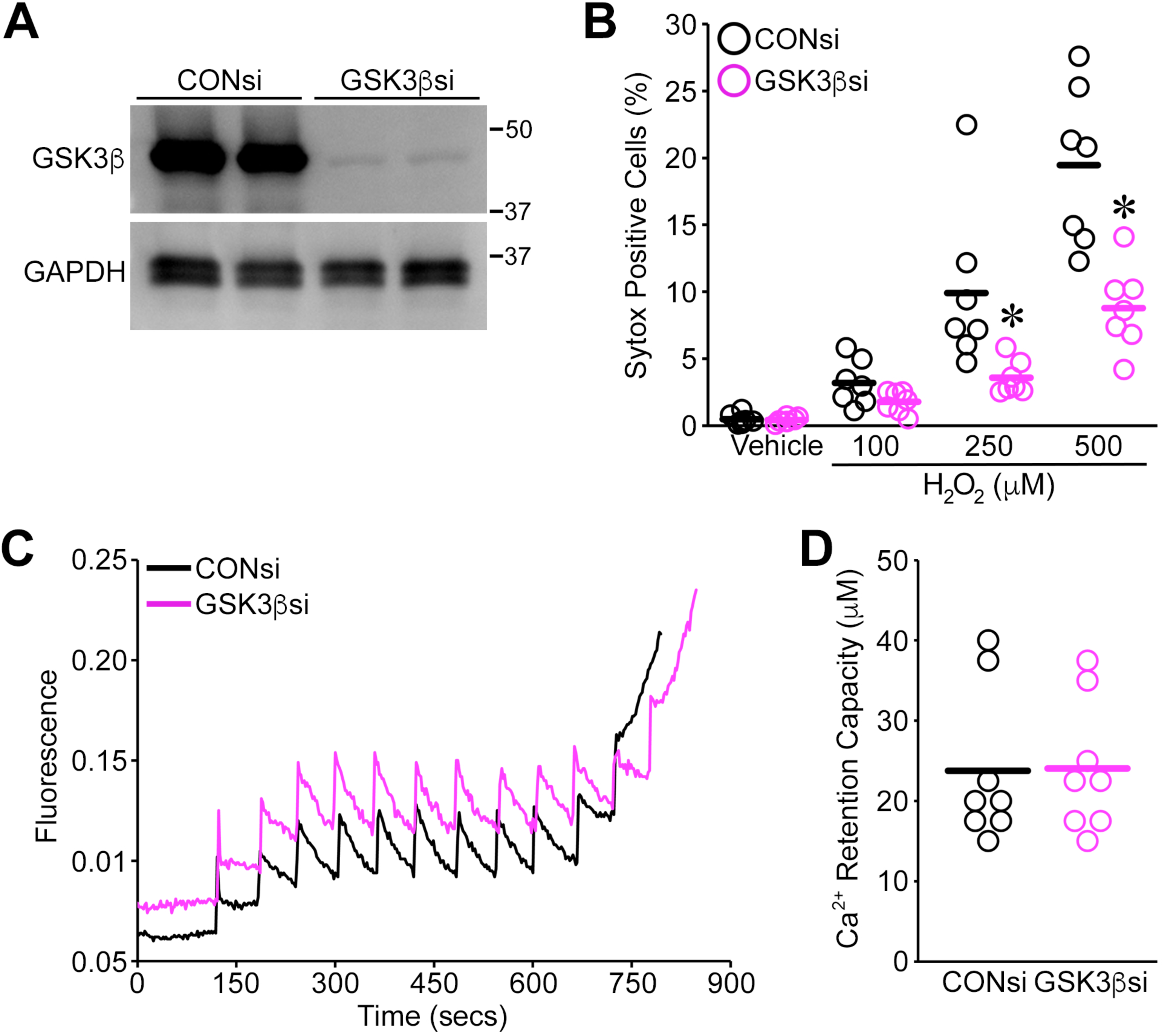
GSK3β silencing protects MEFs against oxidative stress-induced cell death independent of changes in MPT. Wild-type MEFs were transfected with non-targeting (CONsi) or GSK3β-specific (GSK3βsi) siRNAs (100nM) for 48hrs. **A**) Western blotting for GSK3β confirmed depletion of the protein in GSK3βsi-transfected MEFs. GAPDH was used as a loading control. **B**) CONsi and GSK3βsi-transfected MEFs were treated with H_2_O_2_ (100, 250 or 500μM) for 4hrs and cell death was measured by Sytox green staining. **C**) Representative traces of CRC measurements using Calcium Green-5N in digitonin-permeabilized CONsi and GSK3βsi-transfected MEFs. **D**) Quantification of CRC data in the CONsi and GSK3βsi-transfected MEFs. Each individual point represents one independent cell isolate. Bar represents the mean. \**P*<0.05 vs. CONsi.

### GSK3β directly interacts with and phosphorylates CypD

A fraction of the cellular pool of GSK3β has been proposed to reside in the mitochondria matrix [17,25,26,27]. In this regard, mouse hearts were sub-fractionated into mitochondrial and cytosolic fractions and probed for GSK3β. As shown in **Figure 3A**, there was a considerable amount of GSK3β in the mitochondrial fraction. Alkali extraction of mitochondrial membrane proteins confirmed that the GSK3β was in the soluble fractions of the mitochondrion and not integrated into any membrane (**Figure 3B**). The presence of GSK3β in the matrix means that it could interact with and phosphorylate CypD, and therefore directly regulate MPT. To test whether the two proteins could interact, we incubated His-tagged CypD with increasing amounts of kinase-active GST-GSK3β and then pulled the His-CypD down using a Co^2+^-agarose column. Using this approach, we found that GSK3β could directly bind to CypD in a dose-dependent manner (**Figure 3B**). We then repeated the assay in ATP-containing buffer to enable kinase activity and then probed the complexes using antibodies that recognized phosphorylated serine/threonine residues. We found that increasing concentrations of GST-GSK3β resulted in increased serine/threonine phosphorylation of recombinant CypD (**Figure 3B**). These data indicate that GSK3β can indeed bind to and phosphorylate CypD, at least *in vitro*.

**Figure 3:**
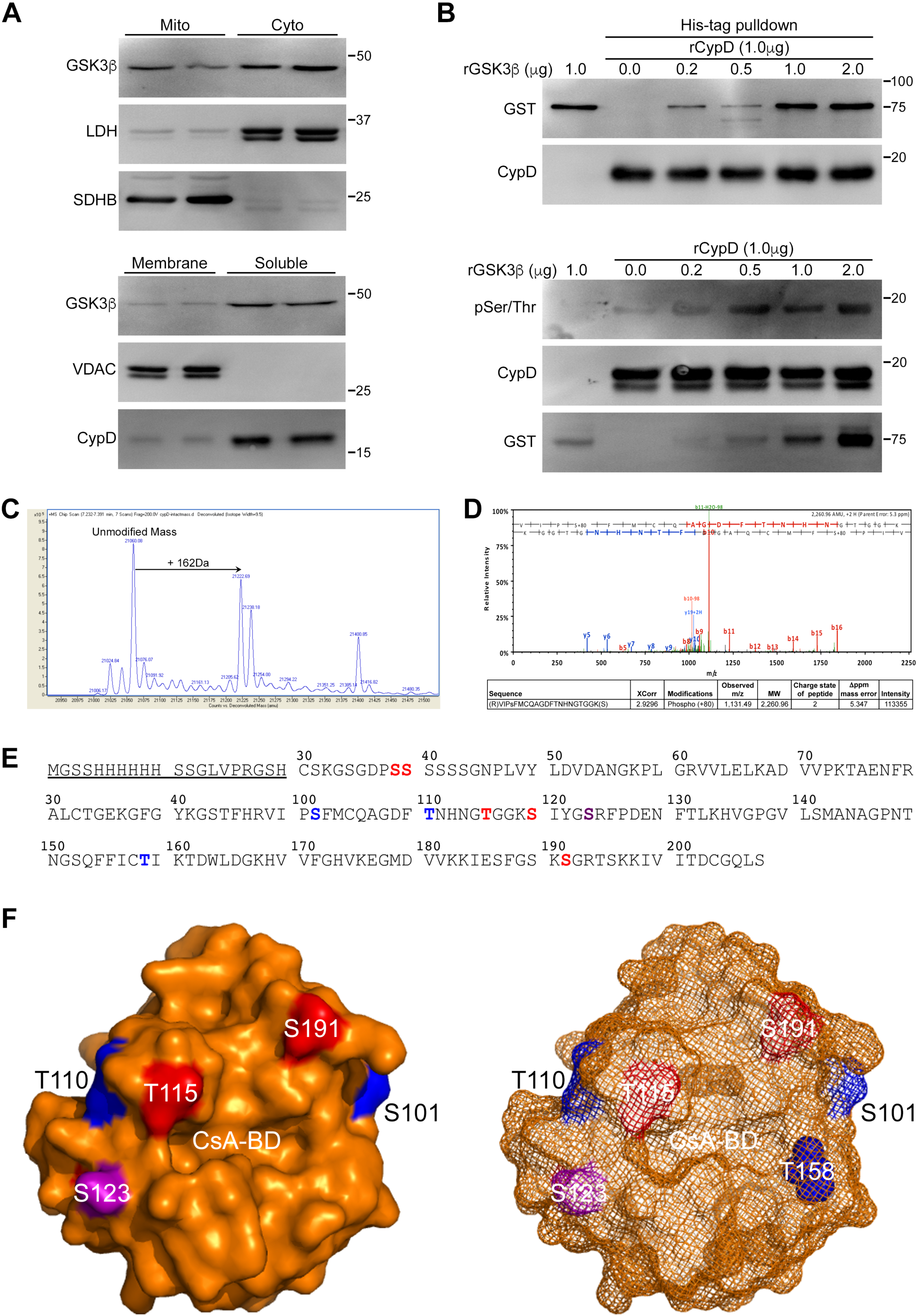
GSK3β localizes to mitochondria and can directly interact with and phosphorylate CypD. **A**) Upper panels, Western blotting for GSK3β, lactate dehydrogenase (LDH, cytosolic), and succinate dehydrogenase (SDHB, mitochondrial) in cardiac mitochondrial and cytosolic subfractions. Lower panels, Western blotting for GSK3β, VDAC (membrane), and CypD (soluble) in alkali extracted membrane and soluble fractions of cardiac mitochondria. **B**) Upper panels, recombinant GST-rat GSK3β was incubated with His-tagged human CypD, purified on Co^2+^-agarose column, and the complexes Western blotted for GST and CypD. Lower panels, recombinant GST-GSK3β was incubated with His-tagged human CypD in kinase buffer supplemented with ATP, purified on Co^2+^-agarose column, and the complexes Western blotted for phosphoserine/threonine, GST and CypD. **C**) Mass analysis of phosphorylated recombinant CypD demonstrating the presence of two phosphorylation sites. **D**) Example of MS-MS spectra indicating S101 as a putative GSK3β phosphorylation site on CypD. **E**) Amino acid sequence of the recombinant His-tagged human CypD depicting canonical GSK3β consensus sites (red), the putative phosphorylation sites identified in our analyses (blue), and a site previously hypothesized to be a key GSK3β phosphorylation residue (purple). **F**) Surface representations of human CypD illustrating the positions of the various putative phosphorylation sites and their proximity to the catalytic CsA-binding domain (CsA-BD).

### Proteomic identification of amino acids on CypD phosphorylated by GSK3β

To identify the specific serine and/or threonine amino acids that were phosphorylated by GSK3β, we purified the His-tagged CypD after the kinase reaction and subjected it to LC-MS/MS. We obtained 20 unique peptides and 26 unique spectra that covered 93% of the protein sequence. **Figure 3C** shows a mass shift of 162 Da, which indicated that CypD was being phosphorylated simultaneously on 2 residues by GSK3β. Three potential phosphopeptides were identified: (R)VIPsFMCQAGDFTNHNGTGGK(S) S101, (R)VIPSFMCQAGDFtNHNGTGGK(S) T110, and (K)HVGPGVLSMANAGPNTNGSQFFICtIK(T) T158. Nonetheless, we found no evidence that the (R)VIPSFMCQAGDFTNHNGTGGK(S) peptide was simultaneously phosphorylated at both S101 and T158, suggesting that the mass shift was due to 2 populations containing S101+T158 or T110+T158. However, none of the sites were unequivocal in their identification. For example, regarding the S101 site the XCorr score was good, mass error was low and fragment coverage was reasonable (**Figure 3D**), but there were no fragment ion matches in this region. We were able to obtain fragment ions for the T110 and T158 peptides, but coverage was limited. That being said there was still enough confidence to pursue them further.

CypD contains several GSK3β consensus S/TxxxS/T^P^ phosphorylation sequences (**Figure 3E**, red and purple amino acids) where the second serine/threonine residue requires pre-phosphorylation by another kinase in order for GSK3β to phosphorylate the first residue [28]. Interestingly, none of the sites we identified were part of a consensus sequence (**Figure 3E**, blue amino acids). Analysis of the CypD PDB structure 3QYU demonstrated that both S101 and T110 were in proximity to the cyclosporine-A binding site, implying that could affect CypD’s isomerase activity (**Figure 3F**). However, S101 is not conserved in the mouse orthologue so we did not pursue this residue further. T158 appears to be internal and therefore may not be accessible to GSK3β in intact mitochondria (**Figure 3F**), but we still wished to follow this site up. In addition, although we did not identify it in our analyses, S123 (**Figure 3E,F**, purple amino acid) has been proposed as a target for mitochondrial GSK3β ◻◻◻◻ so we also investigated whether this site was a regulator of CypD and MPT.

### Putative GSK3β phosphorylation sites do not modulate CypD’s activity or function in MEFs

To determine whether phosphorylation of the T110, T158, or S122 sites influenced CypD and, therefore, MPT and cell death, we generated recombinant GST proteins containing mouse WT-CypD or phosphonegative (alanine) or phosphomimetic (aspartate) mutants at the mouse equivalent sites T109, T157, and S122 (**Figure 4A**). Assessment of peptidyl prolyl isomerase activity of the CypD proteins revealed no difference in enzymatic activity between the mutant and WT proteins (**Figure 4B**). To assess the effects of the phosphosite mutants on CypD’s regulation of MPT, we infected CypD-null MEFs with adenoviruses encoding each CypD construct (**Figure 4C**) and assessed MPT using the CRC assay. As expected, expression of WT-CypD in the CypD-deficient cells greatly reduced the CRC compared to βGal-infected MEFs (**Figure 4D,E**). All of the phosphomutant CypD proteins reduced the CRC to a similar extent as the wild-type protein (**Figure 4D,E**). Finally, we investigated the effects of the mutants on oxidative stress-induced cell death. Re-expression of WT-CypD in the CypD null MEFs significantly increased the amount of Sytox staining in response to 500 μM H_2_O_2_ (**Figure 4F**). Expression of the CypD mutants increased H_2_O_2_-induced cell death to the same extent as the wild-type protein (**Figure 4F**). Taken together, these data indicate that phosphorylation of these residues does not affect the function of CypD and its ability to positively regulate MPT and cell death.

**Figure 4:**
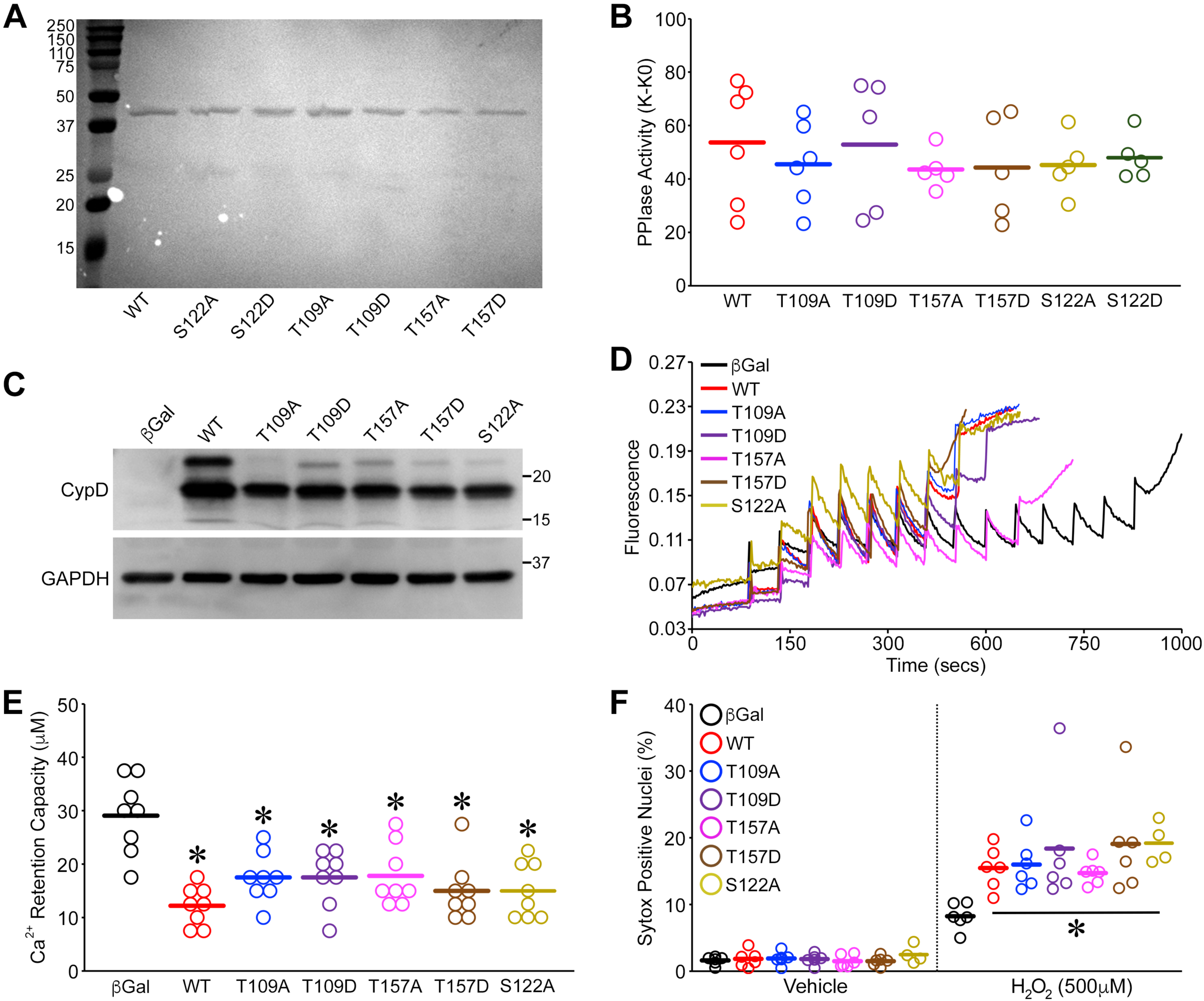
CypD phosphomutations do not alter MPT or oxidative stress-induced death in CypD-null MEFs. **A**) Coomassie staining of purified GST-encoding constructs of wild-type (WT) mouse CypD and CypD mutated at S122, T109 or T157 to alanine (A) or aspartate (D). **B**) PPIase activity of the purified CypD proteins in the presence and absence of the CypD substrate (N-succinyl-Ala-Ala-Pro-Phe) was measured, data fitted in first order rate kinetics to yield rate constants, K and K_0_ respectively, and the difference represented as K-K_0_. **C**) CypD KO MEFs were infected with adenoviruses encoding WT, T109A, T109D, T157A, T157D, and S122A CypD for 48hrs and expression confirmed by Western blotting for CypD. GAPDH was used as a loading control. **D**) Representative traces of CRC measurements using Calcium Green-5N in digitonin-permeabilized WT and phosphomutant CypD-infected CypD KO MEFs. **E**) Quantification of CRC data in the WT and phosphomutant CypD-infected CypD KO MEFs. **F**) WT and phosphomutant CypD-infected CypD KO MEFs were treated with 500μM H_2_O_2_ for 4hrs and cell death was measured by Sytox green staining. Each individual point represents one independent cell isolate. Bar represents the mean. \**P*<0.05 vs. WT (PPIase activity) or βGal (CRC and Sytox).

### Specific targeting of GSK3β to the mitochondrial matrix does not affect MPT or oxidative stress-induced cell death in MEFs

Although the identified GSK3β phosphorylation sites on CypD did not affect MPT or cell death, it was still feasible that mitochondrial GSK3β could still influence CypD and/or MPT indirectly. To test this, we generated WT- and DN-GSK3β adenoviruses with the COXVIII mitochondrial localization sequence to target them to the matrix (miWT-3β and miDN-3β). Equivalent expression of these proteins in MEFs was confirmed by Western blotting (**Figure 5A**), and immunocytochemistry for the C-terminal HA-tag established the correct mitochondrial localization (**Figure 5B**). However, there was no difference in the CRC in either the miWT-3β or miDN-3β expressing MEFs compared to the βGal-infected control cells (**Figure 5C,D**). When we examined sensitivity to oxidative stress-induced cell death, unlike expression of total WT- and DN-3β (**Figure 1B**), neither miWT-3β nor miDN-3β significantly altered the dose-dependent increase in Sytox staining elicited by H_2_O_2_ (**Figure 5E**). Thus, mitochondrial matrix GSK3β is not sufficient to increase MPT sensitivity and cell death.

**Figure 5:**
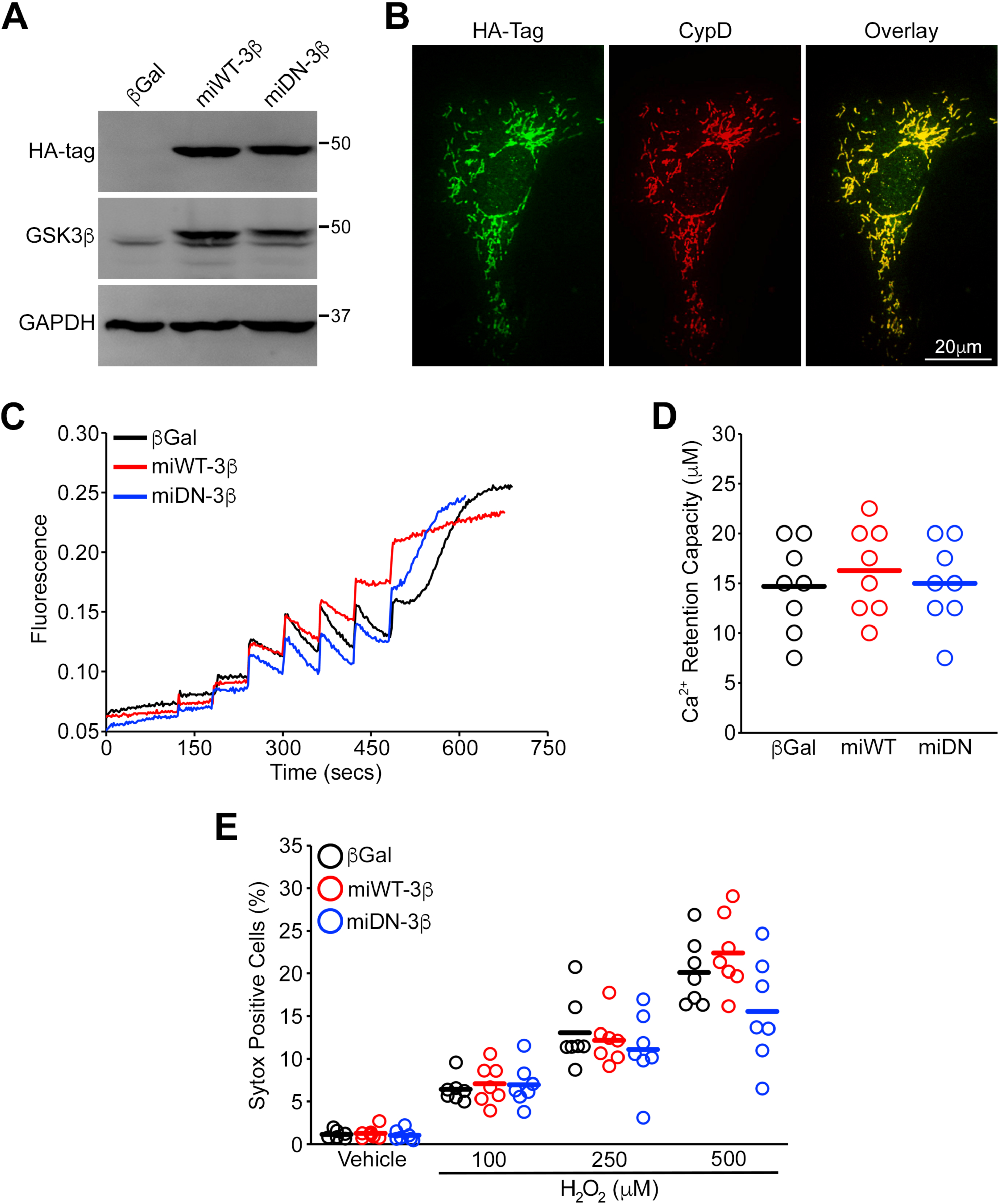
Overexpression of mitochondrially-targeted WT- or DN-GSK3β does not alter MPT or oxidative stress-induced death in MEFs. Wild-type MEFs were infected with adenoviruses encoding βGal (control) or mitochondrially-targeted HA-tagged WT-GSK3β (miWT-3β) or DN-GSK3β (miDN-3β) for 48hrs. **A**) Western blotting for HA and GSK3β confirmed equivalent expression of the two GSK3β proteins. GAPDH was used as a loading control. **B**) Representative images of immunocytochemistry performed on MEFs infected with miWT-3β stained for the HA tag confirmed mitochondrial localization of the protein. Mitochondria were visualized by staining for CypD. **C**) Representative traces of Ca^2+^ retention capacity (CRC) measured using Calcium Green-5N in digitonin-permeabilized βGal, miWT-3β and miDN-3β-infected MEFs. Cells were treated with 2.5μM Ca^2+^ boluses every minute until the peak of fluorescence indicative of MPT was observed. **D**) Quantification of CRC data in the βGal, miWT-3β and miDN-3β-infected MEFs. **E**) βGal, miWT-3β and miDN-3β-infected MEFs were treated with H_2_O_2_ (100, 250 or 500μM) for 4hrs and cell death was measured by Sytox green staining. Each individual point represents one independent cell isolate. Bar represents the mean.

### Mitochondrial GSK3β is not in the matrix and does not interact with CypD in intact MEFs

Our initial data indicated that overexpression of WT-GSK3β could increase MPT Ca^2+^-sensitivity and exacerbate H_2_O_2_-induced cytotoxicity (**Figure 1**). However, our data showing that mutation of putative GSK3β sites in CypD (**Figure 4**) and direct targeting of WT-GSK3β to the mitochondrial matrix (**Figure 5**) suggested that the detrimental effects of GSK3β were indirect. We therefore wanted to re-assess the proposal that GSK3β could localize to the mitochondrial matrix. To this end we isolated cardiac mitochondria and subjected them to increasing concentrations of proteinase K (**Figure 6A**). Using this protocol, there was a progressive digestion of mitochondrial components starting with the outer membrane-associated hexokinase-2 (HK2), followed by the intermembrane space protein apoptosis-inducing factor (AIF), the inner membrane complex-III protein ubiquinol-cytochrome c reductase core protein 2 (UQCRC2). CypD in the matrix was protected at all concentrations of proteinase-K (**Figure 6A**). A significant portion of the mitochondrial GSK3β was digested at the lowest concentration, similar to hexokinase-II (**Figure 6A**), suggesting that the majority is on the cytosolic side of the mitochondrial outer membrane. There were still detectable levels of GSK3β even at the mid-level proteinase-K concentrations, with the digestion profile paralleling that of AIF (**Figure 6A**). However, unlike CypD, GSK3β was completely digested at the higher concentrations of protease. Thus, although GSK3β is present on mitochondria and potentially in the intermembrane space, it is not in the matrix.

**Figure 6:**
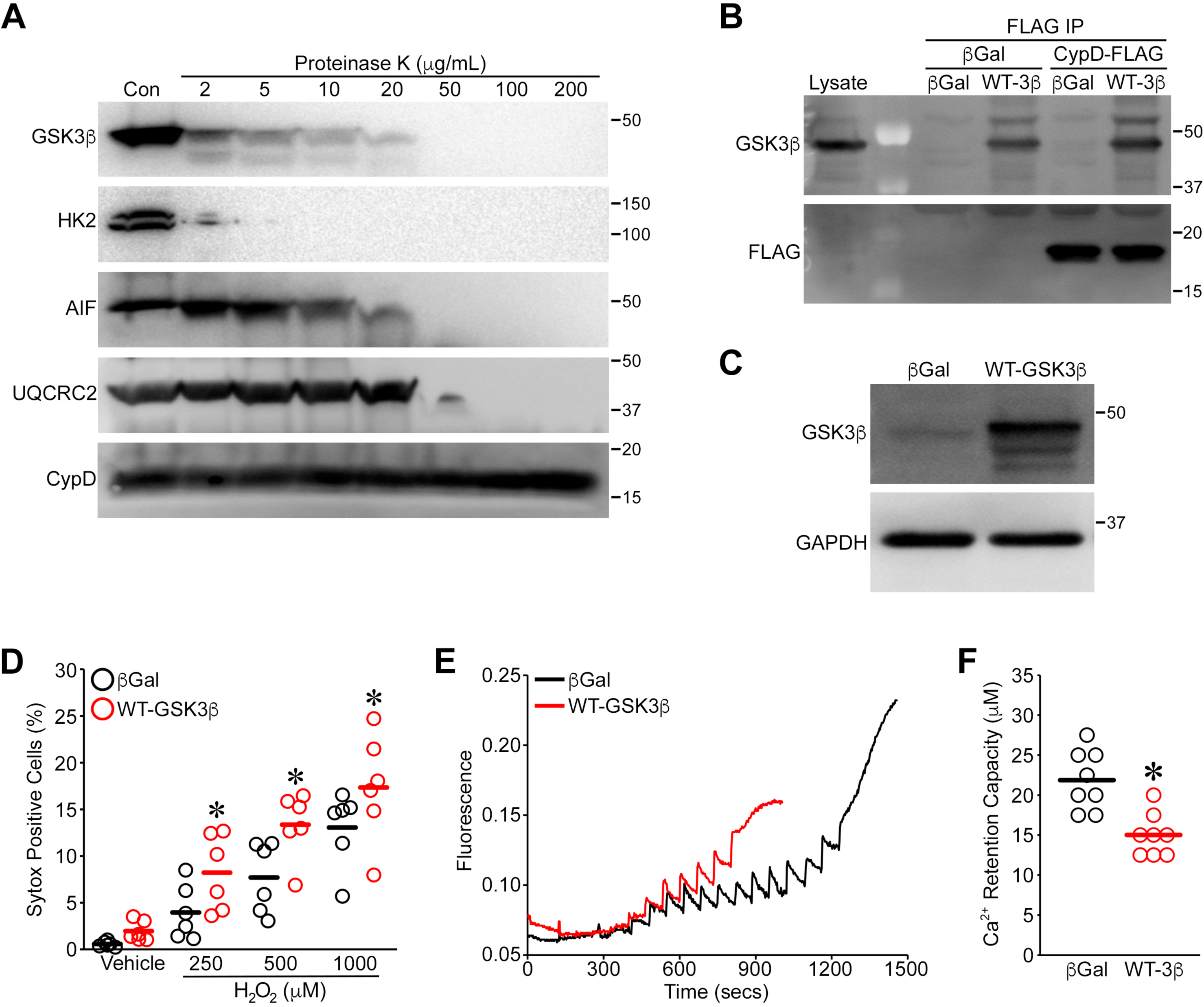
GSK3β is not in the matrix and still sensitizes MEFs to MPT and oxidative stress-induced death in the absence of CypD. A) Cardiac mitochondria were exposed to increasing concentrations of proteinase K and then Western blotted for hexokinase-2 (HK2, mitochondrial surface), apoptosis-inducing factor (intermembrane space), ubiquinol-cytochrome c reductase core protein 2 (UQCRC2, inner membrane), and CypD (matrix). B) MEFs were infected with adenoviruses encoding βGal, WT-GSK3β and/or FLAG-tagged CypD for 48hrs. CypD was immunoprecipitated using an anti-FLAG antibody and the complexes Western blotted for GSK3β and FLAG. C) CypD KO MEFs were infected with adenoviruses encoding βGal, WT-GSK3β for 48hrs. Western blotting for GSK3β confirmed overexpression of GSK3β. GAPDH was used as a loading control. D) βGal and WT-GSK3β-infected CypD KO MEFs were treated with H_2_O_2_ (100, 250 or 500μM) for 4hrs and cell death was measured by Sytox green staining. E) Representative traces of Ca^2+^ retention capacity (CRC) measured using Calcium Green-5N in digitonin-permeabilized βGal, and WT-GSK3β -infected MEFs. Cells were treated with 2.5 μM Ca^2+^ boluses every minute until the peak of fluorescence indicative of MPT was observed. F) Quantification of CRC data in the βGal and WT-GSK3β-infected MEFs. Each individual point represents one independent cell isolate. Bar represents the mean. \**P*<0.05 vs. βGal.

Although our data indicated that recombinant CypD and GSK3β could directly interact *in vitro* (**Figure 3**), our MEF data indicated that this may not be the case in intact cells, especially if they reside in separate mitochondrial compartments. Consequently, we expressed WT-GSK3β and/or FLAG-tagged CypD in MEFs and then immunoprecipitated any resulting complexes using an anti-FLAG antibody. GSK3β did indeed pulldown along with CypD when co-expressed (**Figure 6B**). However, a similar amount of it also precipitated in the βGal-infected cells indicating non-specific binding to the FLAG antibody/bead mix (**Figure 6B**).

### Whole cell expression of GSK3β can increase MPT and cell death in MEFs independent of CypD

Finally, we wanted to test whether the pro-MPT and cell death effects of WT-GSK3β were truly independent of CypD. We therefore overexpressed WT-GSK3β in CypD-null MEFs (**Figure 6C**) and reassessed cell death and MPT. Even in the absence of CypD, WT-GSK3β was still able to enhance H_2_O_2_-induced Sytox positivity (**Figure 6C**), as was seen in wild-type MEFs. Similarly, the previously observed reduction in CRC was recapitulated in the MEFs lacking CypD (**Figure 6E,F**). Thus, any ability of GSK3β to promote MPT and oxidative stress-induced cell death is independent of CypD.

## Discussion

CypD, a core regulator of the MPT, is a potential therapeutic target for the inhibition MPT-related cell death and disease [1,2,3,4,5]. Therefore, the identification of factors such as GSK3β that regulate its activity is critical. In this study, we demonstrated that overexpression of GSK3β sensitized cells to MPT and oxidative stress-induced cell death and GSK3β was able to directly interact with CypD and phosphorylate residues S101, T110 and T158 *in vitro*. However, gain- or loss-of-function mutations of these residues failed to alter CypD’s activity and its ability to positively regulate MPT and oxidative stress-induced cell death. Similarly, direct targeting of GSK3β to the mitochondrial matrix did not affect MPT or cell death. Finally, the cytoprotective effects of genetic GSK3β inhibition were independent of any changes in MPT sensitivity and GSK3β was still able to exacerbate MPT and oxidant cytotoxicity in the absence of CypD. These data indicate that, contrary to previously proposed models, the pro-death function of GSK3β is independent of any physical and/or functional interaction with CypD.

Juhaszova *et al*., demonstrated GSK3β to be a nodal regulator of the MPT machinery, with multiple protective interventions converging to inhibit GSK3β [23]. Subsequent studies have confirmed this ability of GSK3β to modulate MPT and cell death in response to toxic stimuli such as oxidative stress [17,18,23,24]. Yet despite this knowledge, the specific mechanism(s) by which GSK3β promotes MPT are still the subject of debate. Due to several recent reports [17,18,25,26,29], the proposal that GSK3β translocates to and/or resides in the mitochondrial matrix where it promotes MPT through direct binding to and phosphorylation of CypD has gained considerable traction. However, in these studies CypD phosphorylation only correlated with GSK3β activation, and no study to date has teased out the specific sites that GSK3β phosphorylates on CypD and whether this phosphorylation is indeed necessary for the promotion of MPT by GSK3β. Rasola *et al.,* proposed residues S38, S39 and S123 as potential GSK3β phosphorylation sites in human CypD [17]. Other potential consensus sites in human CypD include T115, S119, and S191 (**Figure 3E**). These amino acids are contained in canonical GSK3β consensus sequences of S/TxxxS/T^P^, where phosphorylation of the first S/T residue requires prior phosphorylation of the second S/T by a “priming” kinase [28]. However, our proteomic analyses did not identify any of these sites as targets of GSK3β. Sequence comparison shows that S38 and S39 are not conserved in the mouse. The S123 residue is present in mouse CypD (as S122), but in our study mutation of this amino acid to alanine did not alter CypD’s activity or its ability to facilitate MPT and oxidative stress-induced cell death.

The potential GSK3β phosphorylation sites on human CypD we did identify were S101, T110 as well as T158. Interestingly, none of these residues were contained in an S/TxxxS/T consensus sequence, which was previously considered to be essential for phosphorylation. However, considerable evidence is now emerging that GSK3β can phosphorylate multiple proteins at non-canonical sites [30,31,32]. As S101 is not present in mouse CypD, we focused on T110 and T158 (T109 and T157 in the mouse). Yet, when we mutated these sites to either alanine or aspartate we observed no difference in the activity of CypD. Moreover, reconstitution of these mutants into CypD-deficient cells restored sensitivity to MPT and cell death to a comparable degree as the wild-type protein. Consequently, although GSK3β can directly phosphorylate CypD *in vitro*, such post-translational modifications have zero effect on CypD’s regulation of MPT. This would indicate that the observed ERK-related CypD phosphorylation [17] is in fact GSK3β independent, suggesting that another kinase(s) may be responsible. Multiple serine and threonine amino acids are present on the surface of CypD [33], any of which could be potentially phosphorylated. In this regard, Parks *et al.* reported that S42 in CypD was phosphorylated in mitochondrial calcium uniporter (MCU)-deficient mice [19]. Aspartate mutants of S42 sensitized cells to Ca^2+^-induced pore opening, underscoring other signaling events, which could regulate MPT.

In the present study, we confirmed that overexpression of GSK3β could sensitize cells to Ca^2+^-induced MPT and oxidant-induced death. Yet despite this, our phosphomutant studies suggested that CypD phosphorylation is not in fact required for GSK3β’s ability to positively regulate MPT and cell death. Consistent with this, targeted overexpression of either active or inactive GSK3β in the mitochondrial matrix failed to alter the response of the fibroblasts to MPT-inducing stimuli. While we did find that GSK3β was present in mitochondria, limited protease digestion assays indicated the kinase was not present in the matrix. Thus, although GSK3β can phosphorylate CypD *in vitro*, the fact that it is not in the same submitochondrial compartment as CypD suggests that such an event cannot occur in an intact cell. This was further corroborated by the recapitulation of the exacerbated MPT and cell death response by GSK3β overexpression in MEFs lacking CypD. Taking these data together, we must conclude that any effects of GSK3β on MPT are independent of CypD phosphorylation. This would be consistent with the study of *Clarke et al.*, who found that attenuation of MPT by preconditioning, which increases phosphorylation, and therefore inactivation of GSK3β [23,24], was not associated with changes in mitochondrial protein phosphorylation [34]. It should also be noted that although genetic inhibition of cellular GSK3β did attenuate oxidative stress-induced cytotoxicity, this was dissociated from any changes in MPT sensitivity. This could suggest a possible disconnect with regards to modulation of MPT between the pro-survival effects of GSK3β inhibition vs. the pro-death effects of GSK3β activation. Das *et al.* found that protection of cardiac myocytes by GSK3β inhibition was primarily associated with reduced ATP consumption through reduced adenine nucleotide flux across the OMM, rather than a direct effect on the MPT pore [36]. A very recent study similarly showed that the infarct-sparing effects of novel inhibitors were independent of changes MPT and did not require CypD [37].

Regardless, any capacity of GSK3β to regulate MPT and MPT-dependent cell death must be through indirect mechanisms. Studies have reported that inhibition of GSK3β reduces the interaction of CypD with ANT [24,26]. However, the fact that GSK3β’s effects do not require CypD suggests that disruption of the ANT-CypD complex is not at play here. That being said, our data showing that GSK3β is present in the IMS implies that it could still potentially physically and/or functionally interact with ANT on the cytosolic side of the IMM. Several groups have reported an interaction between ANT and GSK3β [24,35], but whether this contributes to the pro-MPT actions of this kinase has not been tested. We observed the majority of the mitochondrially-associated GSK3β was on the OMM as most of it was digested at the lowest concentration proteinase alongside HK2. Indeed, GSK3β-induced dissociation of HK2 from VDAC1 has been correlated with enhanced MPT-induced cell death [38,39]. Regulation of the pro-apoptotic VDAC2 isoform has also been put forward as a mechanism [36,40]. However, others have failed to find an interaction between GSK3β and VDAC [29], and depletion of any or all of the three VDAC isoforms does not directly affect MPT responsivity [9,10]. Lastly, GSK3β phosphorylates anti-apoptotic member of the Bcl 2 family of proteins, Mcl-1, thereby targeting Mcl-1 to ubiquitin-dependent degradation [41]. This is intriguing as cardiac-specific deletion of Mcl-1 evoked a cardiomyopathy that could be rescued by deletion of CypD [42] and down-regulation of Mcl-1 triggered MPT in leukemia cells [43]. The exact mechanism(s) by which Mcl-1 regulates the MPT pore and whether its degradation plays a role in GSK3β’s pro-MPT and cell death effects are the subjects of our current investigations.

In conclusion, our study demonstrates that while GSK3β can sensitize cells to MPT and oxidative stress-induced death, these effects are independent of phosphorylation of CypD. Moreover, inhibition of GSK3β, while cytoprotective, is not associated with any changes in MPT sensitivity. Together, these findings disprove the hypothesis that direct phosphorylation of CypD is necessary for the facilitation of MPT and MPT-dependent cell death by GSK3β.

## Acknowledgements

This work was supported by National Institutes of Health grants HL094404 (to C.P.B.). The content is solely the responsibility of the authors and does not necessarily represent the official views of the National Institutes of Health.

